# COMPUTATIONAL STUDIES OF CARGO TRANSPORT THROUGH THE NUCLEAR PORE COMPLEX

**DOI:** 10.64898/2026.02.23.707554

**Authors:** Sanjeev K. Gautam, Rozita Laghaei, Afshin Eskandari Nasrabad, Rob D. Coalson

## Abstract

Nuclear Pore Complexes (NPCs) are large protein complexes in eukaryotic cells that span the double-membrane of the nucleus and regulate bi-directional transport between nucleus and cytoplasm. T h e NPC core is lined by intrinsically disordered protein chains called nucleoporins (Nups) which form a selective barrier where large macromolecules (cargoes) need to bind to nuclear transport receptors (NTRs) such as Karyopherins (Kaps) to cross. Previous experimental results have suggested that not only Nups but Kaps, too, are important in the transport process of other NTRs/NTR-cargo complexes. In this work, we assess the role of Kaps in the transport of other NTRs (specifically, NTF2s) through the NPC, a process referred to as the “Kap-centric transport model”. Here, using coarse-grained MD simulation we show that Kaps are able to direct NTF2s into the Nup meshwork, which leads to their increased flow. Our results also suggest that NTRs follow specific lanes inside the pore to maximize efficient transport.

## Introduction

The nuclear pore complex (NPC) is a large protein complex that spans the double-membrane of the nucleus in eukaryotic cells and regulates bi-directional transport between nucleus and cytoplasm. This includes transport of RNA and ribosomal proteins moving from the nucleus to the cytoplasm, as well as proteins, carbohydrates, signaling molecules and lipids moving into the nucleus.^1, 2^ The NPC is a key component of nucleocytoplasmic transport and regulatory processes such as gene regulation and signaling. The dysregulation of NPC function is implicated in several diseases, ranging from various forms of cancer to autoimmune diseases.^3, 4^

The NPC channel is roughly in the shape of an hourglass spanning the nuclear envelope, with an inner-ring diameter that varies between ∼45 nm in the constricted state and up to ∼70 nm when dilated, as reported in recent cryo-ET and structural studies,^5^ and is lined by multiple copies of ca. 10- intrinsically disordered protein chains called nucleoporins (Nups).^6–8^ These nucleoporin chains are rich in hydrophobic phenylalanine–glycine (FG) repeats, which can interact in various ways depending on their sequence composition. While FG–FG contacts are proposed in some models to contribute to the formation of a selective barrier, their exact role remains under active discussion and likely varies across different classes of FG-Nups. ^6, 9–11^ Depending on the cargo type, transport events take from several milliseconds to hundreds of milliseconds with a typical flux of hundreds of molecules per second per NPC.^1, 7^ Small inert molecules generally diffuse passively through the NPC, with permeability gradually decreasing as molecular size increases rather than showing a strict cutoff near 50 kDa, as observed in previous studies.^12, 13^ Larger macromolecules, however, typically require recognition by nuclear transport receptors (NTRs) such as Karyopherins (Kaps), which bind cargo and facilitate translocation through the FG-Nup barrier.^14^ This selective mechanism, which prevents non-specific large biomolecules from passing through the NPC, is essential for maintaining proper cellular function.

The NPC allows bidirectional transport of cargo-NTR complexes with very high transport rates. However, the precise mechanism of this transport process is under debate. Based on several *in vitro* and *in silico* studies over the years, a number of transport models have been proposed. Depending on the major role of FG-Nups or Kaps, these models can be broadly characterized as “FG-centric” (virtual gate/polymer brush model,^15^ selective phase model/hydrogel model,^16^ forest model^9^) and “Kap-centric” (reduction of dimensionality,^17^ reversible collapse,^18^ molecular velcro^19^). FG-centric models assert that FG-Nups are solely responsible for establishing the selective barrier. This also implies that Kaps are only responsible for transporting cargoes and do not play a role in forming the selectivity barrier. T h e Virtual Gate model proposes that FG-Nups create an entropic barrier where the energy gained by binding of transport receptors to FG repeats assists in paying the entropic penalty to cross the pore. In the selective phase model, cohesive hydrophobic interactions between FG repeats create a sieve-like meshwork that allows small molecules to cross through the mesh while blocking large non-specific macromolecules. The forest model proposes that individual Nups can adopt an extended or collapsed conformation depending on their size, composition and physiological properties, which results in multiple transport pathways within the pore. In contrast, the Kap-centric transport model proposes a dual role of Kaps where they not only transport cargoes but also assist in transport of other NTR-cargo complexes.^19–21^ This is said to be brought about by the presence of two distinct population of Kaps i.e. a “slow-phase” Kap population that is responsible for reshaping the FG-mesh by avid multivalent binding to FG-repeats and a “fast-phase” Kap population is responsible for transporting cargoes across the pore. Slow phase Kaps are bound strongly (low *K_D_ <* 1*µ*M, where K_D_ is the equilibrium dissociation constant indicating binding affinity) to FG-repeats at steady state leading to a saturation of binding sites which allows fast-phase Kaps (high *K_D_ >* 10*µ*M) to move quickly through the pore.^19, 20^ The two phases of Karyopherins were first revealed by Kapinos et al.^20^, who studied the binding kinetics of karyopherin*β*1 (Kap*β*1) with surface-grafted FG-Nup assemblies. The presence of slow phase Kaps bound to FG-Nups also resulted in fast movement of NTF2s near the center of the pore, presumably due to reduction of dimensionality.^22^ Recently, Fragasso et al. found experimental evidence for two phases of Kaps in biomimetic nanopores formed by tethering Nsp1 chains to the inner wall of a solid-state nanopore.^23^

Although there has been experimental evidence to support both types of models, no consensus on a universally applicable model for nucleo-cytoplasmic transport (NCT) has been formed. The heterogeneous nature of the pore, fast simultaneous transport of multiple NTRs (10^3^ molecules per second per pore)^24^ combined with limitations in imaging techniques makes it challenging to spatiotemporally resolve this process.^8, 24, 25^ Artificial mimics and computational models of NPC have been useful in capturing the nature of NCT; however, to our knowledge, there have been no studies focused on the effect of Kaps on the transport of other NTRs under pore confinement.

Previous computational studies have examined nucleocytoplasmic transport using coarse-grained representations of FG-Nups and transport receptors, highlighting the role of FG-Nup density, crowding, and self-regulation in sustaining efficient transport. ^26–28^ These models demonstrate how spatially heterogeneous FG-Nup organization gives rise to effective transport barriers and robust flux under crowded conditions. Building on this framework, we focus here on how such density organization gives rise to spatially resolved transport pathways and Kap-centric transport lanes within the pore.

In this work, we employ molecular dynamics simulations to study the transport of two well characterized NTRs, Nuclear Transport Factor 2 (NTF2) and Importin-*β* (Imp-*β*) through a coarse-grained, 1-bead per amino acid model of NPC. The NPC core is constructed by grafting 32 copies of 5 different types of FG-Nup chains in an eight-fold symmetry following the work by Winogradoff et al.,^29^ while NTRs are modelled as rigid bodies with binding sites that interact with FG patches. We aim to re-evaluate the Kap-centric transport model and the effect of Kaps (Imp-*β*) on NTF2 transport through the pore. Our simulations support some aspects of Kap-centric model while there are also some discrepancies. Notably, with the force field utilized in this work, we do not find a stable strongly bound population of Kaps (slow phase) in the pore. Also, there is no swelling of FG-mesh upon binding by Kaps as previously observed in experimental measurements.^19–21^ Our data suggests that Kaps occupy the center of the pore, and from this location are able to direct NTR traffic into the FG Nup mesh. Also, we observe several distinct pathways or “lanes” that NTF2s follow to cross the pore with flux being highest through the high FG density region. Based on our simulation data we propose that Kaps are responsible for pushing NTRs into the Nup mesh where the attractive interactions between FG repeats and binding sites on NTRs help expedite the transport process.

## Methods

### CG model for unfolded proteins

All coarse-grained MD simulations were performed using implicit solvent one bead per amino acid (1-BPA) model developed by Onck et al.^30, 31^ with recently updated cation-pi interactions.^32^ In this model, each amino acid is represented as a single bead placed at the *C_α_* position and has a mass of 120 Da. The bond separating the beads is modelled as a stiff harmonic potential with an equilibrium distance of b=0.38 nm: and *K_b_* = 8038 kJ nm*^−^*^2^ mol*^−^*^1^.

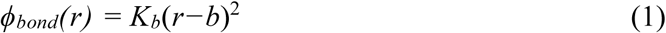

The backbone potentials, i.e., bending and torsion, are assigned via an explicit mapping of Ramachandran data of a library of the coil regions of proteins that distinguishes flexible, stiff, and regular amino acids. Non-bonded interactions between amino acids consist of hydrophobic, charged, and cation-pi interactions. T h e relative hydrophobicities of amino acid residues are normalized between 0 to 1 based on a partition energy measurement, i.e., the free energy of transfer between polar and apolar solvents. The hydrophobic interaction potential has the following form:

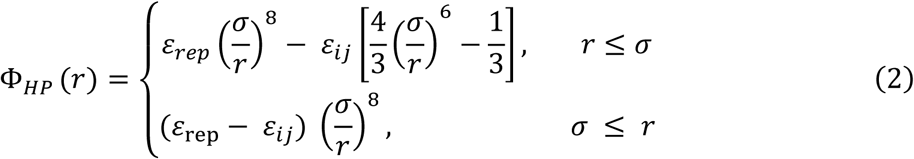

Here, *ε_rep_* = 10 kJ/mol and 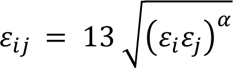 with *ɛ_i_* being the hydrophobicity value of amino acid i normalized between 0 to 1 and *α* = 0.27 is an empirical scaling factor. The interaction potential also includes pairwise electrostatic interactions described by a modified Coulombic law:

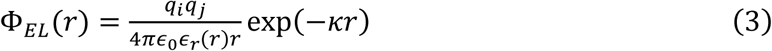

where *κ* = 1.0 *nm^−^*^1^ is the Debye screening coefficient accounting for salt concentration in cytoplasm,^33^ while *ɛ_r_*(*r*) is a distance dependent dielectric constant of the form:

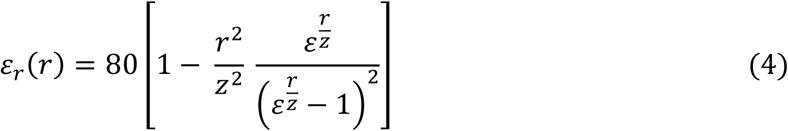

For a residue pair where a positively charged residue interacts with a residue with an aromatic side chain, instead of the hydrophobic potential above a cation-pi potential of the form:

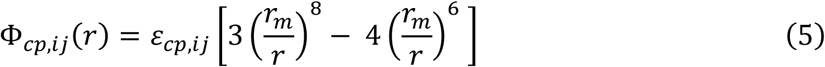

is invoked, where the equilibrium distance is *r_m_* = 0.45 nm and *ɛ_cp,ij_* is the pairwise interaction parameter introduced by Jafarinia et. al, representing the attractive strength between aromatic (F, Y and W) and positively charged residues (R and K).^32^

### CG model of the NPC

To create the NPC core, we graft 32 copies of 5 different types of nucleoporin chains (FG-Nups) in an eight-fold symmetry, similar to the work of Winogradoff et al.^29^ These chains, Nup145N, Nic96, Nsp1, Nup49, Nup57, consist of 732, 139, 467, 245 and 77 amino acids respectively, with 32x5=160 chains in total (Fig. 1). Each amino acid is represented by a coarse- grained spherical bead of diameter 0.60 nm and, as noted above, is assigned a mass of 120 Da. In general, FG motifs commonly appear in the FG-Nup sequence either alone or in “patches” such as FGFG, FxFG, or GLFG combinations. Therefore, FG patches in these chains were identified by checking through the amino acid sequence for F1 (phenylalanine) followed by a G1 (glycine) residue. These residues were then assigned F1 and G1 to differentiate them from other F and G residues that are not in an FG patch.

**Figure 1:**
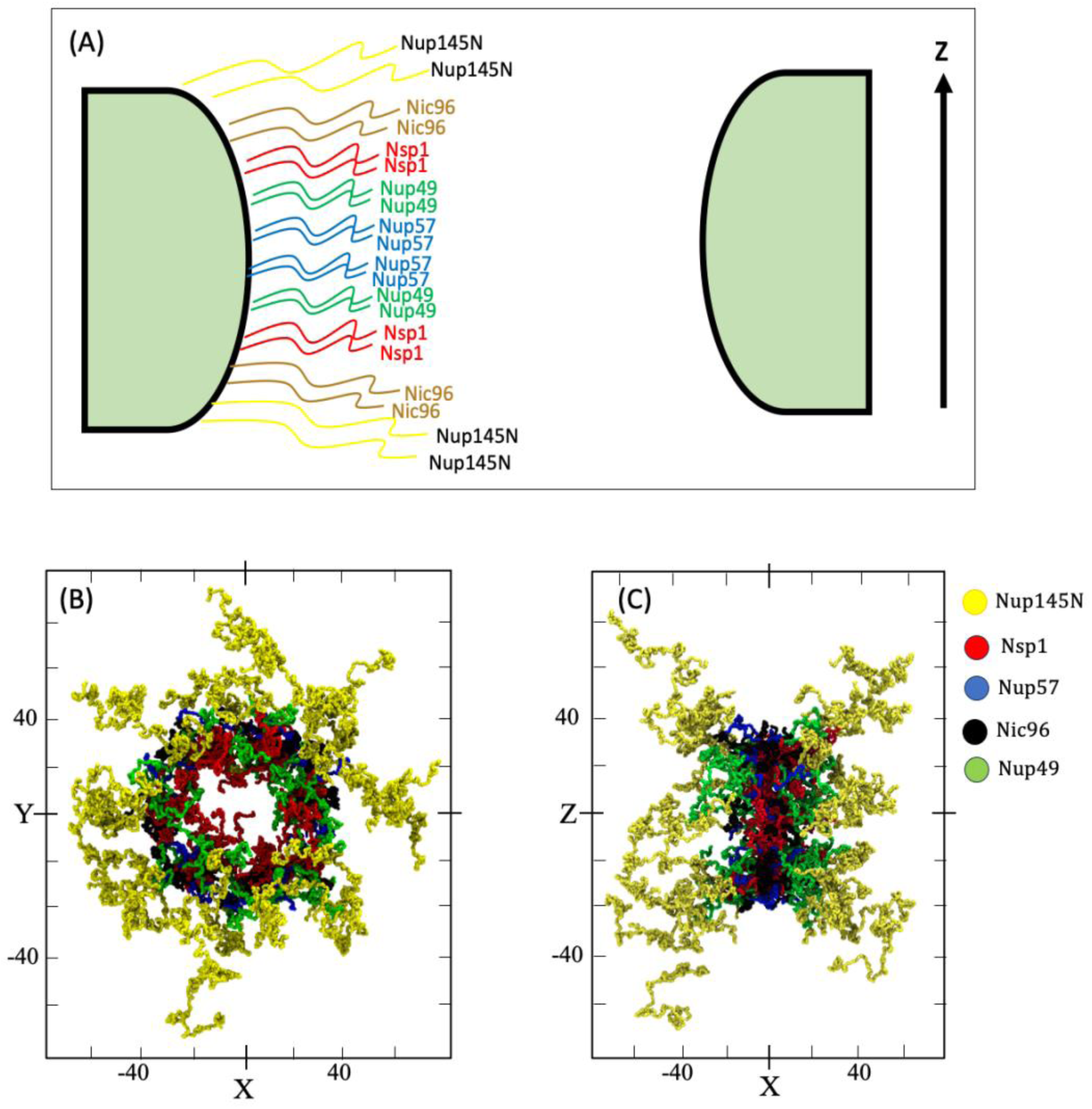
(A) A sketch of the system showing grafted nucleoporin chains of different types. (B) Top view (x–y plane) snapshot of the equilibrated FG-Nup assembly, highlighting the radial organization and overlap of different Nup types within the pore cross-section. (C) Side view (x–z plane) snapshot of the same system, illustrating the axial extent of the FG-Nup network and its confinement within the pore geometry. Axis units are in Angstroms (Å) and with colors denoting different FG-Nup species.

### CG model of transport protein (Importin-***β*** (Kap) and NTF2)

The CG transport molecules were constructed from all-atom crystallographic structure by placing a bead at the *C_α_* position of each amino acid (Fig. 2). Each transport molecule is treated as a rigid body, with its constituent beads fixed in their relative positions throughout the simulation. This means that each timestep the total force and torque on each rigid body is computed as the sum of the forces and torques on its constituent particles. The composition of binding sites in each protein molecule were located based on the work by Isgro and Schulten.^34, 35^ There are 10 binding sites in Importin-*β* (PDB ID: 2QNA) and 6 in NTF2 (PDB ID: 1OUN). If the binding site on the transport molecule contains any of the following residues: R, K, F, Y, W, we impose a binding site interaction with F1 and G1 amino acids in the nucleoporin chains with an experimentally determined interaction strength of *ɛ_BS−F_*_1*G*1_ = 13.75 kJ/mol.^23^ Binding-site locations and corresponding residue lists are given in Appendix Figures A1–A2 and Tables A1–A2.

**Figure 2:**
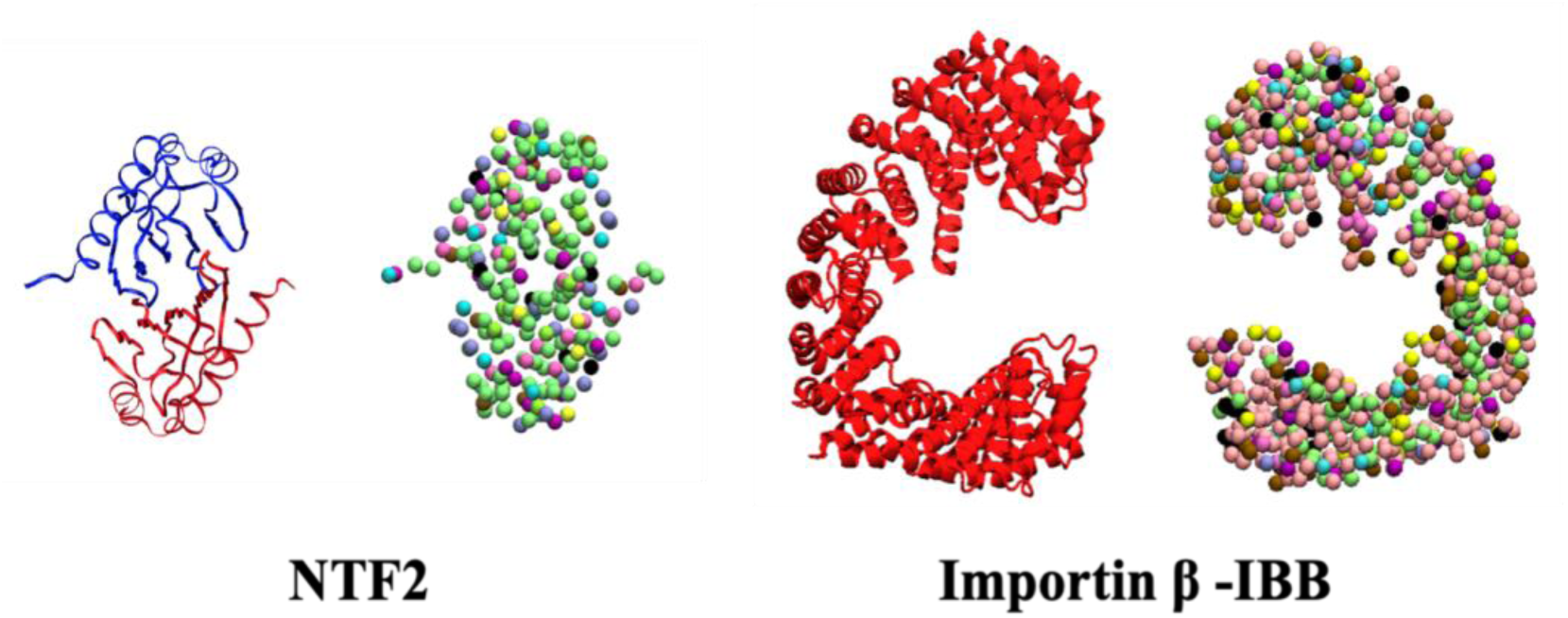
Coarse grained structure of the transport proteins: NTF2 (Left) and Importin-β (right).

### CG simulations of the NPC model

All CG MD simulations were performed using the LAMMPS (Large-scale Atomic/Molecular Massively Parallel Simulator) MD simulation package^36^ with Langevin dynamics integrator at a temperature of 300K. Timestep of 0.03ps and a Langevin friction coefficient of 50 *ps^−^*^1^ (following the work of Ghavami et. al. and other similar studies ^23, 31^) were applied in every simulation with non-bonded interaction cut off distance being 1.5nm. Initially, the cytoplasm (Fig. 3) is filled with NTRs. After an initial equilibration period within the cytoplasmic side, they are allowed to pass through the pore to the right side of the nucleoplasm. The walls of the simulation box are periodic along all three Cartesian directions. In concert with these Periodic Boundary Conditions, we employ a nudging force (f*_z_* ) in +z direction at the box edges which circulates NTRs from the nucleoplasm back into the cytoplasm.^37^ This nudging force is important in maintaining unequal concentrations of NTRs on the two sides of the pore by recycling NTRs from nucleoplasm back to cytoplasm and in turn producing a net flow of NTRs.

**Figure 3:**
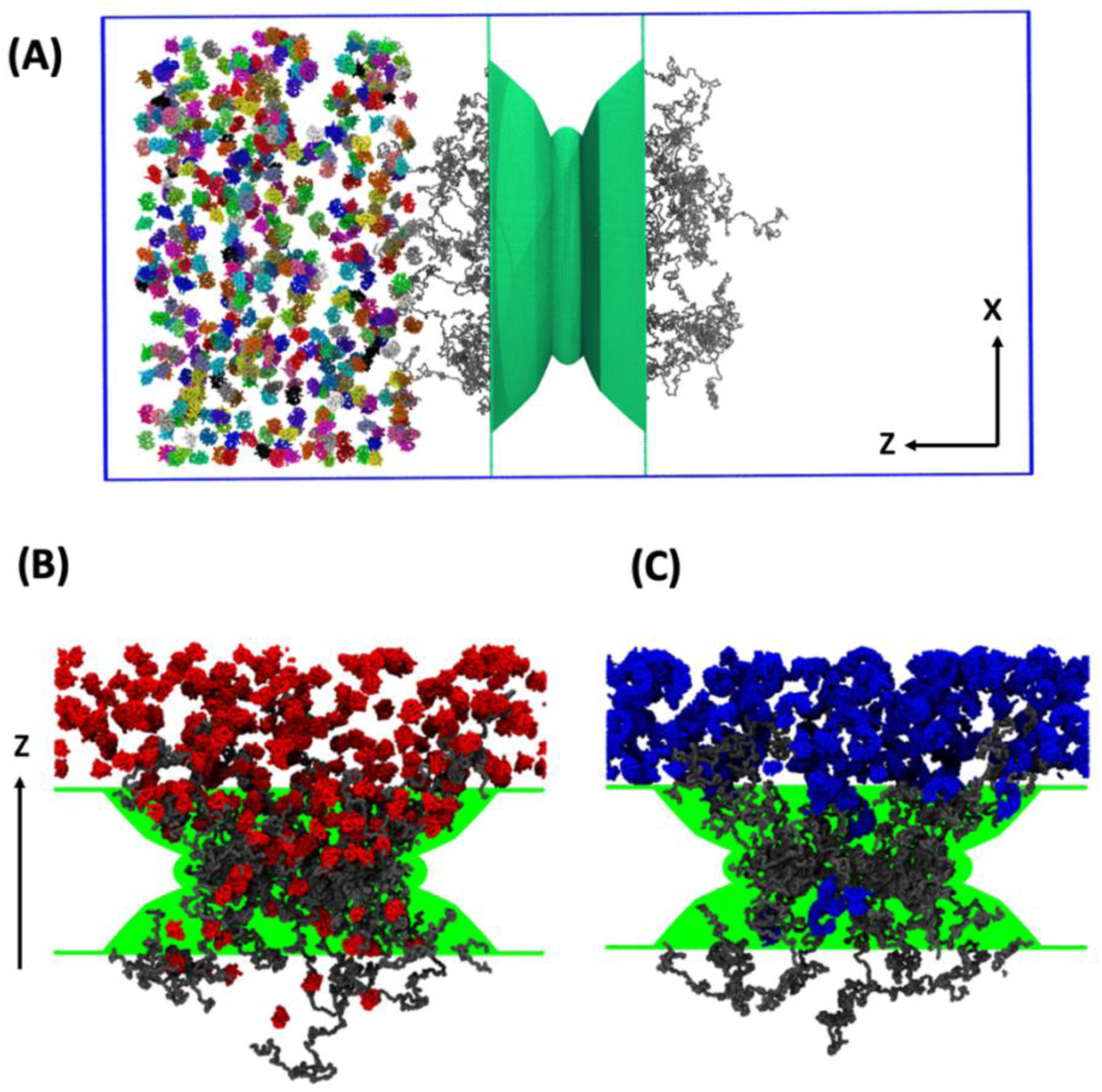
(A) Snapshot of the simulation setup showing 500 NTF2 molecules initially distributed on the cytoplasmic side of the Nuclear Pore Complex (NPC). The coarse-grained FG-Nup assembly forming the pore is shown at the center, separating cytoplasm (left) and nucleoplasm (right). (B) Representative snapshot illustrating NTF2 molecules infiltrating the pore and interacting with the FG-Nup mesh within the central constriction. (C) Representative snapshot showing karyopherins (Kaps) infiltrating the pore, preferentially occupying the central region and interacting with the FG-Nup network.

To study the effect of Kaps on NTF2, we constructed four different simulation setups listed in Table 1. Here, setup C and D are included to observe this effect in the absence of NTF2 binding site interaction with F1, G1 residues.

**Table 1:** List of the simulation setups.

| Setup | NTRs in the setup |
| --- | --- |
| A | NTF2s ($N_{NTF2} = 500$) and Kaps ($N_{ImpB} = 250$); |
| B | Only NTF2s ($N_{NTF2} = 500$), no Kaps ($N_{ImpB} = 0$) |
| C | NTF2s ($N_{NTF2} = 500$) with Kaps ($N_{ImpB} = 250$) and no NTF2 binding site interaction with F1, G1 residues (no NTF2 -F1G1 interaction) |
| D | NTF2s ($N_{NTF2} = 500$), no Kaps ($N_{ImpB} = 0$) and no NTF2 -F1G1 interaction |

## Results

### Assessment of fast/slow phase Kaps

The main evidence for the two phases of Kaps reported in previous studies is the fast transit time for some (fast phase) Kaps and slow transit time for other, strongly bound (slow phase) Kaps.^20, 22, 23^ In our study, when Kaps, otherwise designated as Importin-*β*s (Imp-*β*), enter the pore via the cytoplasmic side, they mostly pass through the pore quickly without strongly binding to the Nup chains. That is, we do not observe a slow phase Kap population. Snapshots extracted from the simulations after 4μs were utilized to compute the radial density profile (DP) of the Nup monomers, NTF2 and Kap molecules in the central region (−3nm < z < 3nm, Fig. 6A) of the pore using the following normalization condition:

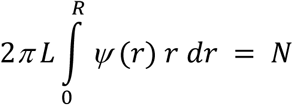

**Figure 4:**
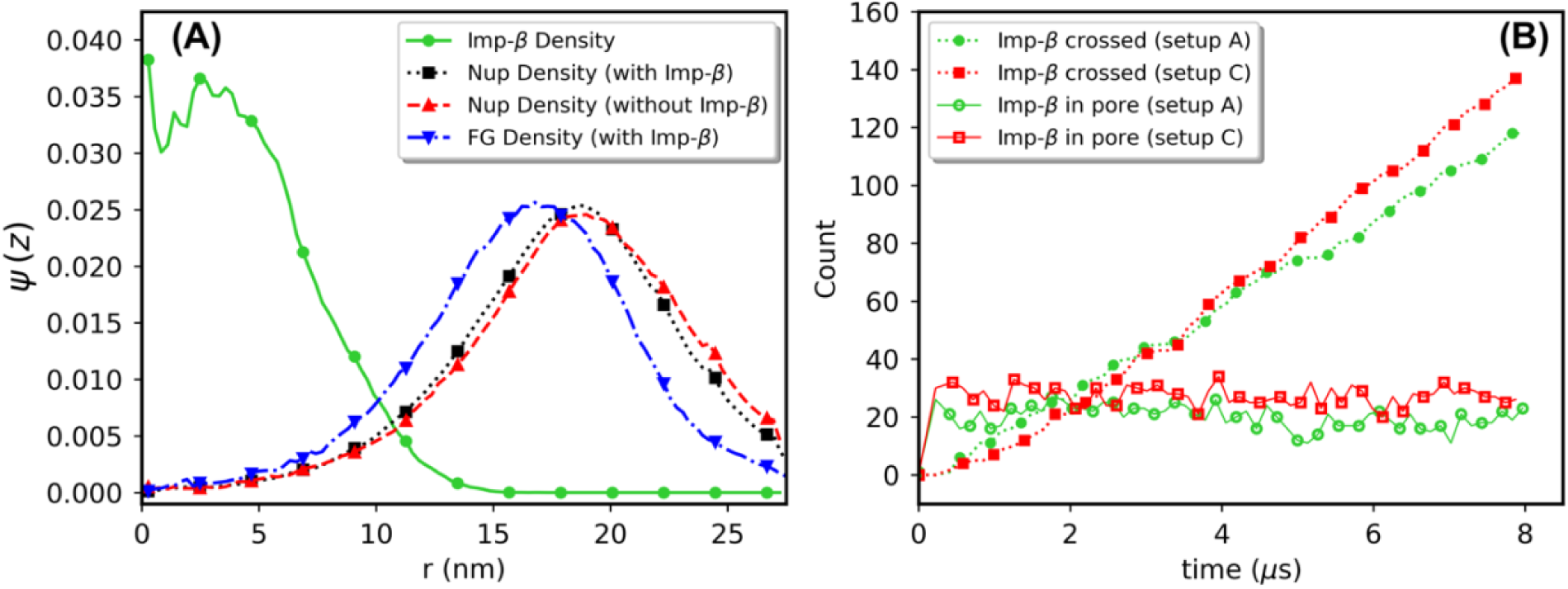
Density profiles of Nup monomers and Kaps for the system with only Kaps (no NTF2s). B: Cumulative Kap count that crossed through the pore and number of Kaps in the pore as a function of time.

**Figure 5:**
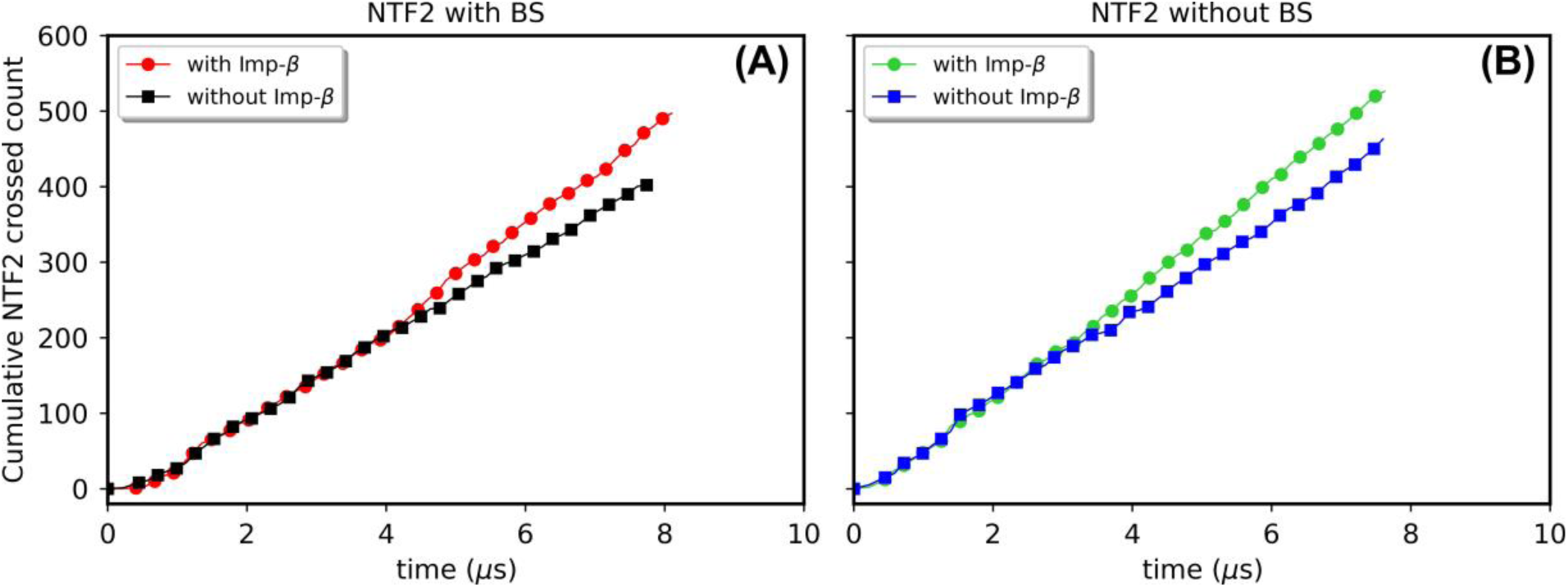
NTF2s crossing the pore versus time for setups A and B (left) and setups C and D (right). “BS” denotes binding sites. The curves within each panel illustrate the effect of Kaps on NTF2 transport under identical binding-site conditions. In both panels, the presence of Kaps increases the slope of the cumulative count curves, indicating an enhanced NTF2 flux. Panels A and B should not be compared directly; instead, the appropriate comparison is between the two curves within each panel.

**Figure 6:**
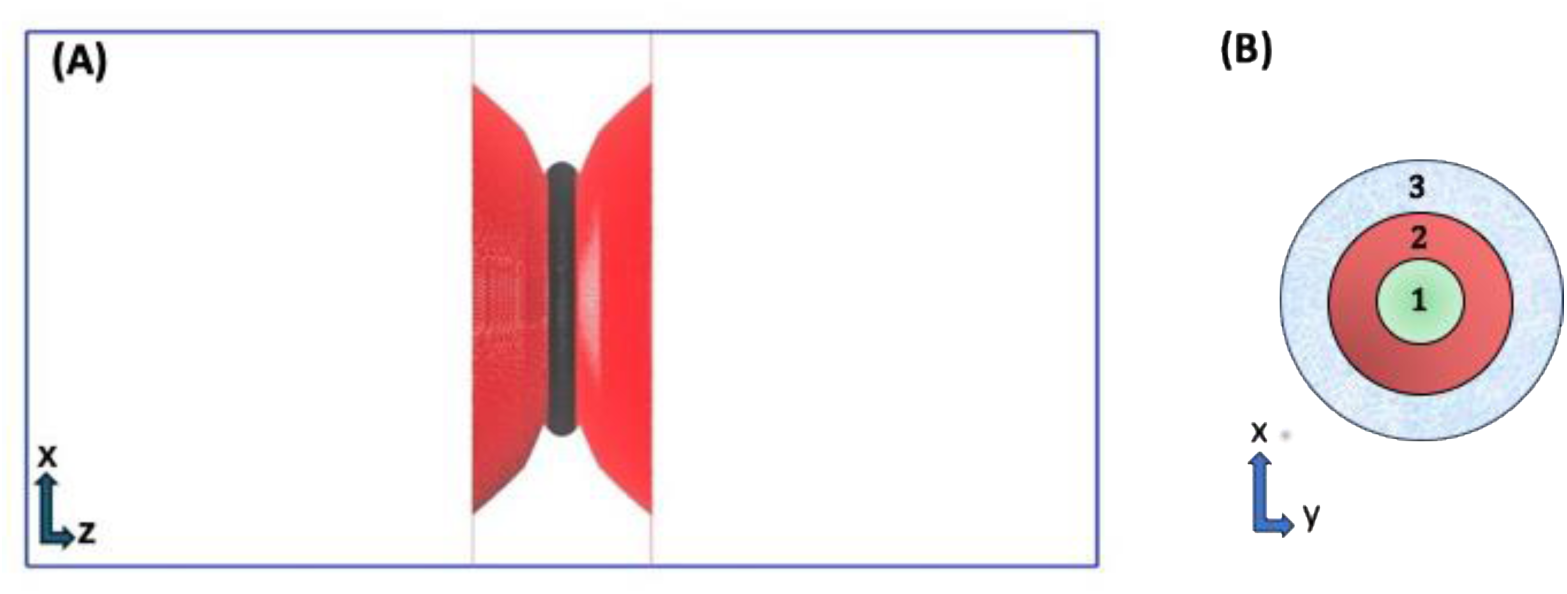
A: Sketch of the central section of the pore shown in black. B: Three different layers in the pore.

where *L* is the length of the central region, *r* is the distance of a monomer/center of mass of a molecule from the cylinder axis, N is the average number of monomers/molecules in the central region, and *ψ (r)* is the number density of the monomer/molecule in a region at a distance r from the center of the pore.

Fig. 4A shows the density profiles of the Nup monomers and Kaps in the central section of the pore. Here, r = 27.5 nm is the grafting surface radius and r = 0 is the center of the pore. The Kap density is highest close to the center of the pore and the presence of Kaps in the pore has minimal effect on the Nup density profile. The total number of Kaps that pass through the pore as a function of time is illustrated in Fig. 4B, indicating the presence of fast-phase Kaps. Additionally, the number of Kaps in the pore over time is displayed, showing approximately *∼* 8% occupancy at steady state.

### Role of Kaps in NTR transport

To investigate the role of Kaps in the transport of other NTRs, we incorporated NTF2s into the cytoplasmic side and analyzed their movement with and without Kaps. To evaluate this effect without the influence of binding sites (BS), we also repeated these simulations without FG binding sites on the NTF2s (setup C, D).

We measured the flux of NTF2 through the pore by counting the number of NTF2 molecules that were recycled from the nucleoplasm back to cytoplasm. Our simulations revealed that a greater number of NTF2 molecules crossed the pore in the presence of Kaps. This finding is illustrated in Figure 5, where the cumulative count of NTF2 molecules crossing the pore is higher when Kaps are included in the system. This trend persists even in the absence of FG-Nup-NTF2 binding site interactions, as shown in panel B of Figure 5. These results confirm that Kaps enhance the flow of NTF2 through the pore, leading to a higher flux.

### Multiple transport pathways or “lanes” in the pore

To investigate the mechanism by which Kaps influence NTF2 flow, we divided the “central section” of the pore (indicated in black in Fig. 6A) into three distinct radial layers. These layers are illustrated in Fig. 6B. Layer 1, located at the center of the pore, is nearly devoid of FG-Nups (cf. Fig. 4A). Layer 2 represents a high-density region of FG-Nups, while Layer 3 is situated close to the grafting surface. We focus on the central section of the pore because it represents the narrowest geometric cross-section and the region where NTF2 transport is most sensitive to local FG-Nup density and Kap occupancy. This region therefore provides the most relevant location for quantifying density distributions, flux, and pathway selection, without implying the existence of a discrete physical bottleneck for NTF2 transport.

Next, we calculated the number of NTF2s that enter the central section of the pore and determined the corresponding layers they pass through. This was done by placing an imaginary dividing surface at the cytoplasmic side of the central section of the pore and measuring the radial distance from the center of each NTF2 molecule entering this section. Only NTF2s that successfully crossed the pore were included in this analysis. Figure 7 categorizes the NTF2s that enter the pore in different layers as a function of time. The results indicate that the number of NTF2s that enter the pore through layer 2 is significantly enhanced by the presence of Kaps. Notably, this trend is also seen when there is no attractive interaction between NTF2s and FG-Nups i.e. setup C vs D. In setup A, the high NTF2 count in Layer 2 is due to a combined effect of Kaps and F1G1-NTF2 interactions, while in setup C, it’s due to Kaps pushing NTF2s into layer 2. These results clearly indicate that Kaps located mostly at the center of the pore, can direct movement of other transport proteins within the pore.

**Figure 7:**
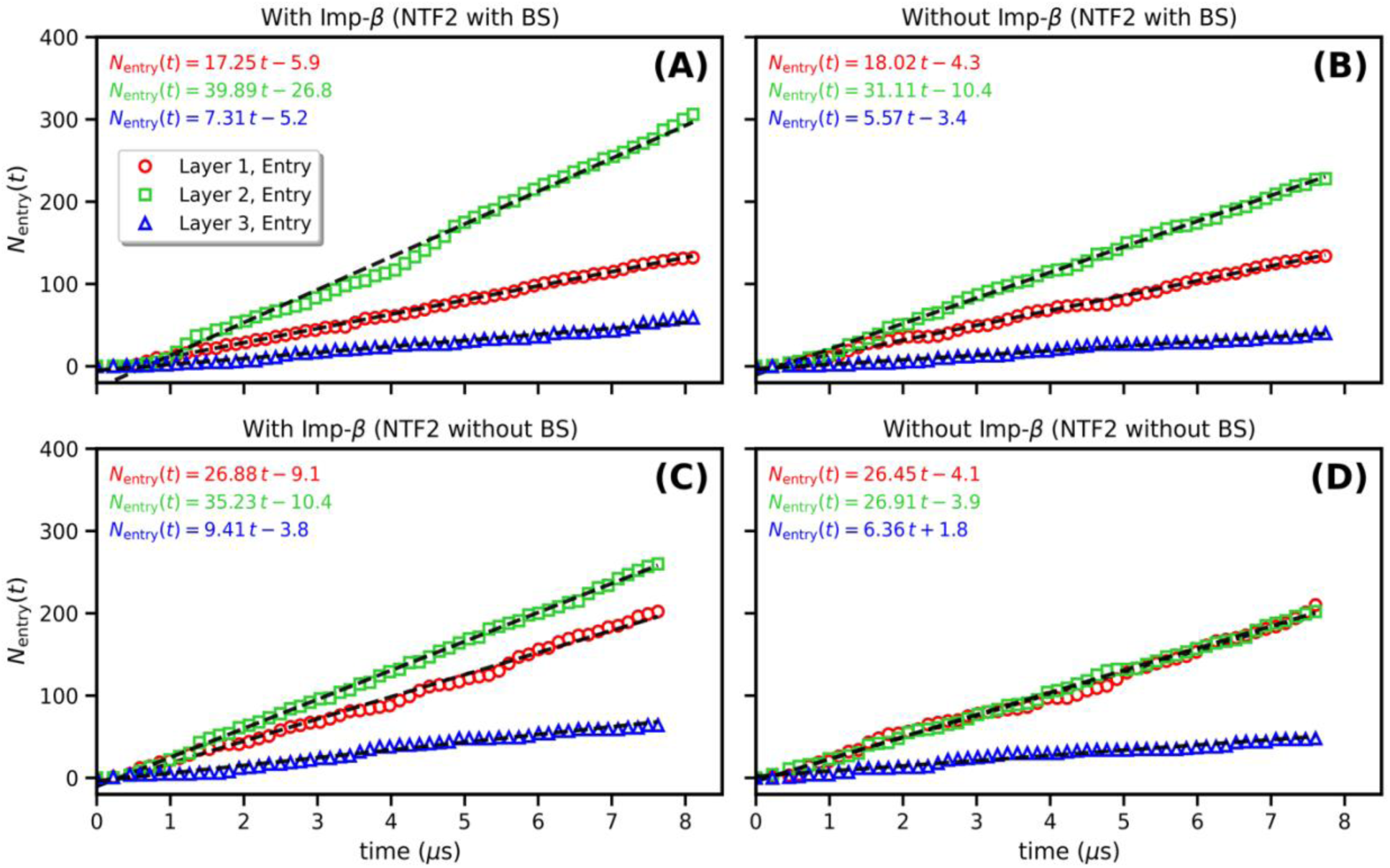
NTF2 entry count (N_entry_) through different layers in the central section of the pore. Here, layer 1 contains the center of the pore, layer 2 is high FG density region and layer 3 is close to the wall. Dashed black lines indicate linear best-fit trends used to quantify entry fluxes, and the corresponding fit equations are shown on each panel.

Figure 8 shows the layer through which NTF2s exit to the right side. In setups A and B, NTF2s primarily exit via layer 2. In contrast, in setups C and D, the absence of F1G1-BS attractive interactions combined with the low Nup density of layer 1 leads NTF2s to prefer exiting through this layer. A comparison of the plots in Figures 7 and 8 indicates that not all NTF2s exit from the layer they initially entered.

**Figure 8:**
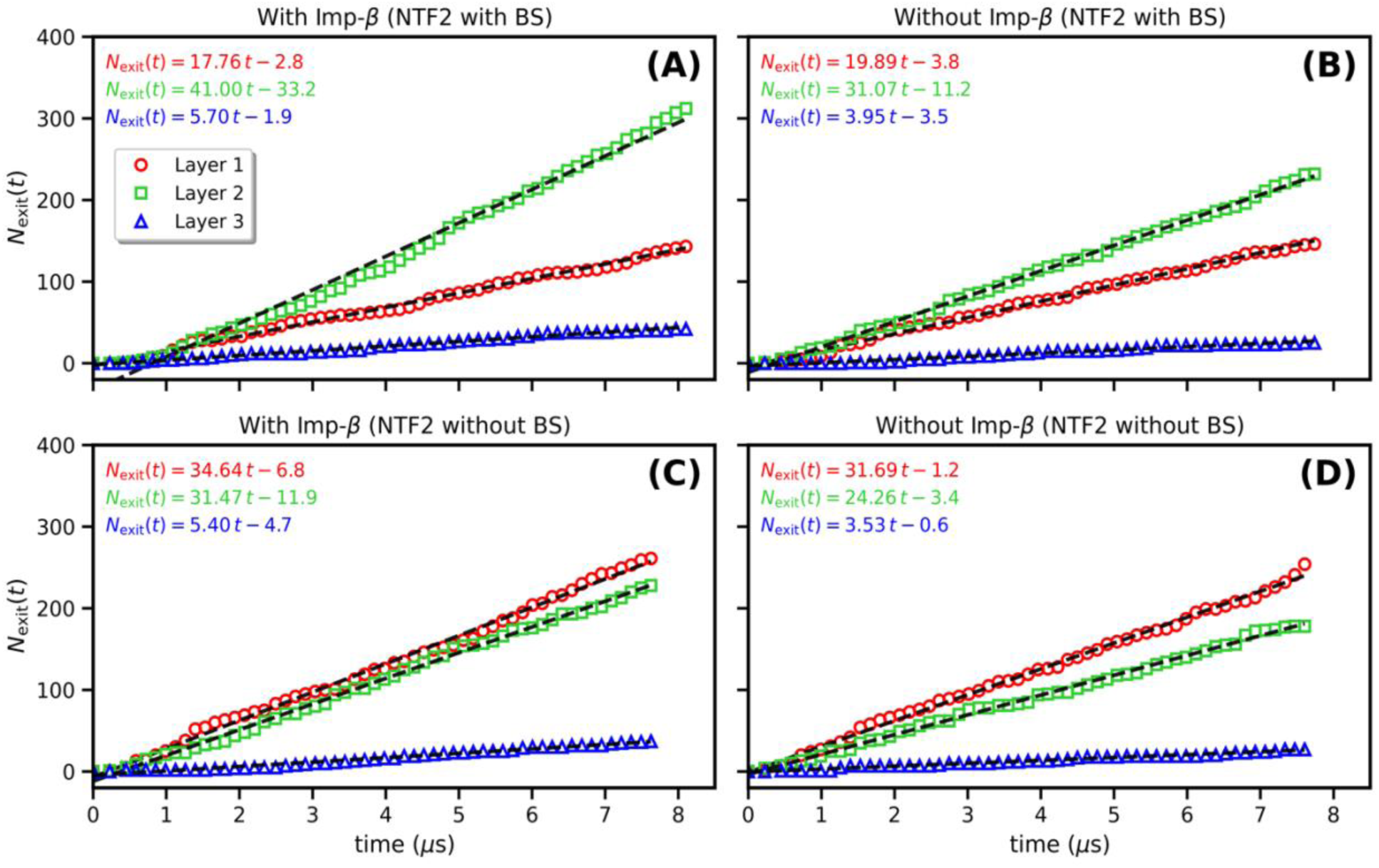
NTF2 exit count (N_exit_) through different layers in the central section of the pore. Dashed black lines indicate linear best-fit trends used to quantify entry fluxes, and the corresponding fit equations are shown on each panel.

### Important role of FG-Nups (Layer 2)

Previous studies have documented the role of FG-Nups in aiding the transport of nuclear transport receptors (NTRs).^33, 38, 39^ When comparing transport through the high FG density layer (Layer 2) across all four setups, it is evident that the highest flux occurs when there is an attraction between F1G1 and NTF2 binding sites, along with the presence of Kaps. In contrast, the flow through the high Nup density layer is lowest when both Kaps and the F1G1-NTF2 interaction are absent.

Figure 9A and B illustrate the number of NTF2s that exit through layer 2 across all four simulation setups. Interestingly, the flux through layer 2 mirrors the trends observed in the overall flux plots (Fig. 5) and the points where the flux value (slope of the NTF2 crossing count vs. time plot) changes also align well. This suggests that the change in flux is primarily due to the increased flow of NTF2s through the high FG density region (Layer 2). The change observed after approximately 4.2 µs in the overall flux in setup A (Fig. 5A) corresponds to an absolute increase of ∼13.8 NTF2 translocation events. This value is comparable to the flux change observed in Fig. 9A (∼14.5 NTF2 translocation events). Similarly, the flux difference between setups C and D at approximately 3 µs corresponds to absolute increases of ∼6.3 NTF2 translocation events for the overall flux and ∼4.4 NTF2 translocation events for the flux through layer 2. Therefore, we conclude that the high flux in setup A is the result of the combined effects of (i) the attractive interaction between F1G1s and NTF2s and (ii) Kaps directing the NTF2s. In contrast, the high flux in setup C is solely attributed to Kaps directing NTF2s to layer 2. Notably, there is no significant difference in the NTF2 flow through layer 1 (Fig. 9C) whether Kaps are present or absent. This also indicates that Kaps do not necessarily block NTF2s as they move through the pore in layer 1 (cf. Fig. 4B).

**Figure 9:**
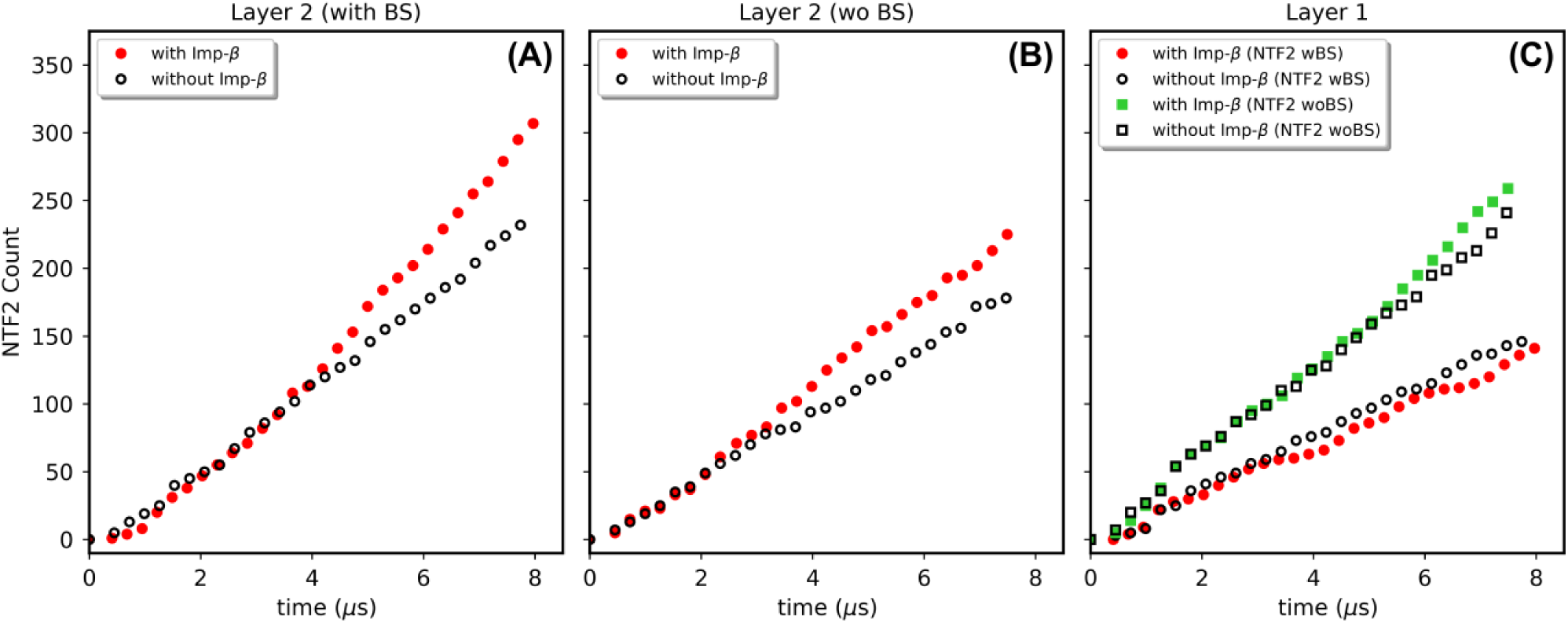
Panel A and B: Flux through the high FG density region (layer 2). Panel C: Flux through Layer 1.

Although Kaps are pushing the NTF2s into the FG mesh as they enter the central section of the pore, without an attractive interaction between F1G1 and NTF2s, the Nups are not able to retain the NTF2s, resulting in their transfer back to Layer 1. Fig. 10 shows a plot of *N_exit_ – N_entry_* vs time where *N_exit_* is the number of NTF2s that exit through a particular layer and *N_entry_* is the number of NTF2s that enter through that layer in the central region of the pore. In setup A and B, due to the presence of binding sites on NTF2s, they tend to stay in the layer in which they originally entered from; however, a small portion of NTF2s in setup B do move from the FG mesh layers (layers 2 and 3) to layer 1. In setup C and D, a large number of NTF2s that entered the pore through layers 2 and 3 exit through layer 1.

**Figure 10:**
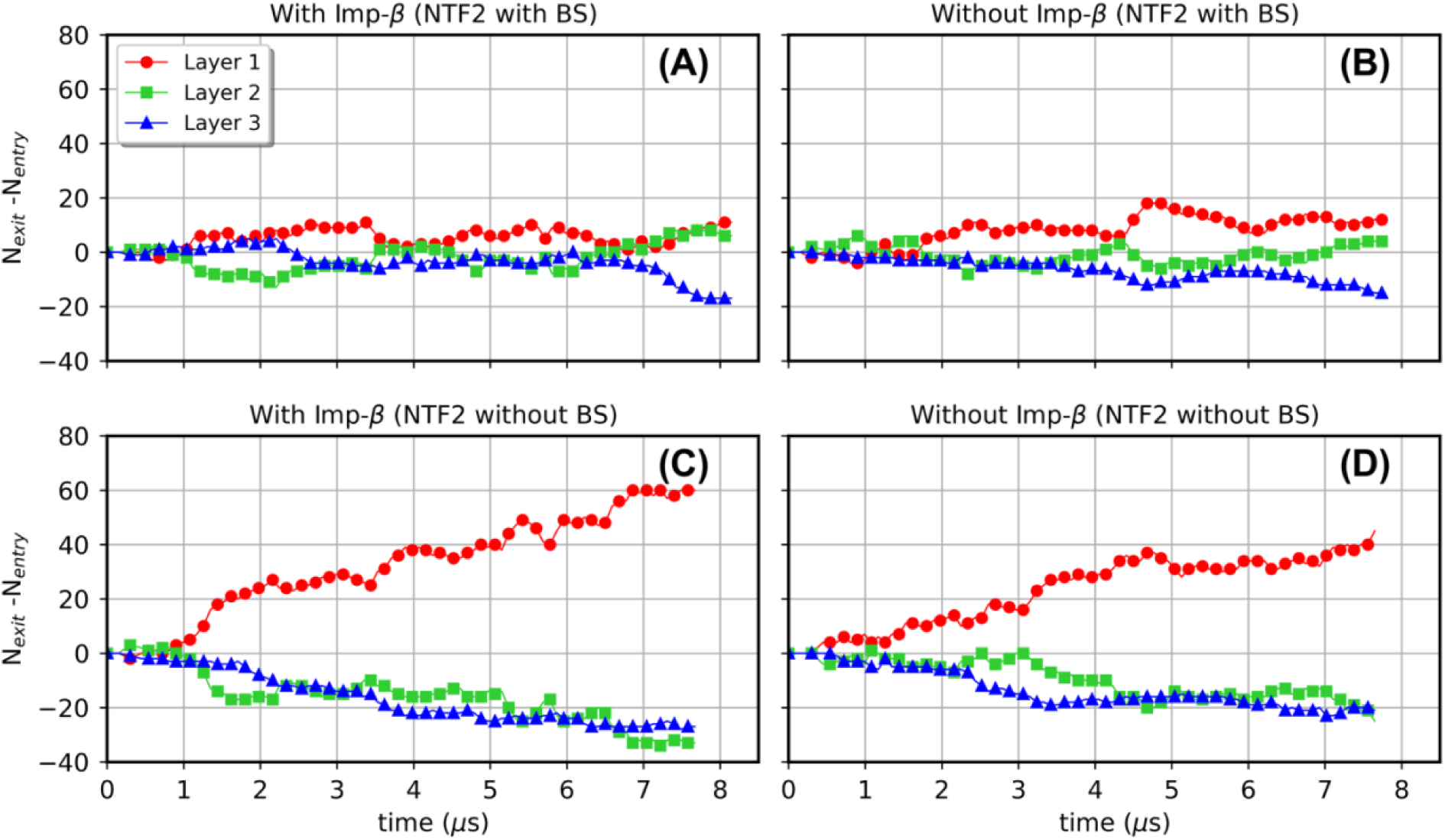
N_exit_ − N_entry_ vs time for the four setups. Here, N_exit_ and N_entry_ is the number of NTF2s that exit and enter the central section of the pore, respectively.

The presence of Importin-β leads to a systematic but modest shift in *N_exit_ – N_entry_,* indicating that Kap-mediated effects on this quantity are secondary in magnitude. This shift does not correspond to a major redistribution of NTF2 trajectories; rather, it reflects a subtle bias in how NTF2s are guided within the pore. Despite its small magnitude, this bias consistently favors transport through the high FG-density region (Layer 2), suggesting that Kaps promote NTF2 entry into regions where FG interactions and local Kap occupancy facilitate more efficient passage.

Taken together, these results indicate that the “lanes model,” which assumes largely layer-preserving transport, is valid for setups A and B. In contrast, in setups C and D the model breaks down, as NTF2s frequently transition from the FG-rich layers to the pore center prior to exit.

NTF2s having attractive interactions with FG-Nups remain mostly in Layer 2, while those without binding sites do not stay in a specific layer but instead move between different layers. Consequently, the lanes model does not seem to apply to NTF2s lacking binding sites, which are instead driven by entropic factors.

### Crossing times

Fig. 11 shows the crossing time of Kaps and their average distance from the pore center. The trend here confirms that Kaps mainly traverse close to the center of the pore and that there is no evidence of slow phase Kaps, which would be characterized by much larger crossing times than the ones observed. In our simulations, we also find that when the binding sites are removed from the NTF2s, they traverse faster through the NPC (setup C and D). Fig. 12 shows a comparison of the times of NTF2 crossing events in setup A vs C. Fast transport and high total flux in setup C can be attributed to the small size of NTF2 molecules (*∼*28 kDa), which is much lower than the 40 - 60 kDa cutoff for diffusion-driven passive transport. Without binding sites, NTF2s behave like small, non-specific molecules (e.g., Thioredoxin (12 kDa), *α*-lactalbumin (14 kDa), GFP (25 kDa)) that can diffuse through the pore without engaging in hydrophobic interactions with FG-Nups. This passive diffusion explains the enhanced speed and highest overall flux observed in setup C.

**Figure 11:**
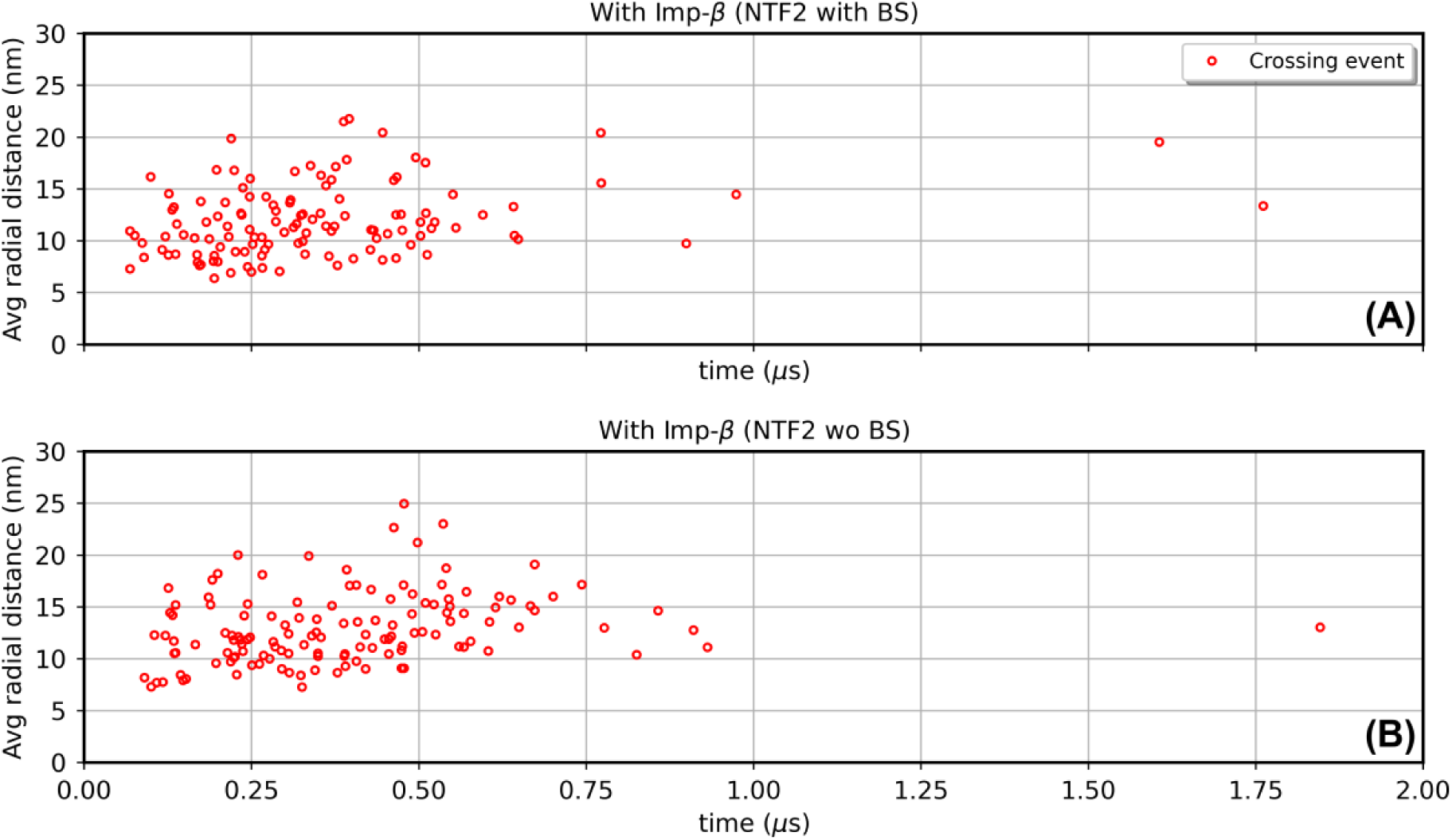
Average radial distance in the pore vs time it takes for the Importin-β to cross the pore for setup A (panel A) and setup C (panel B).

**Figure 12:**
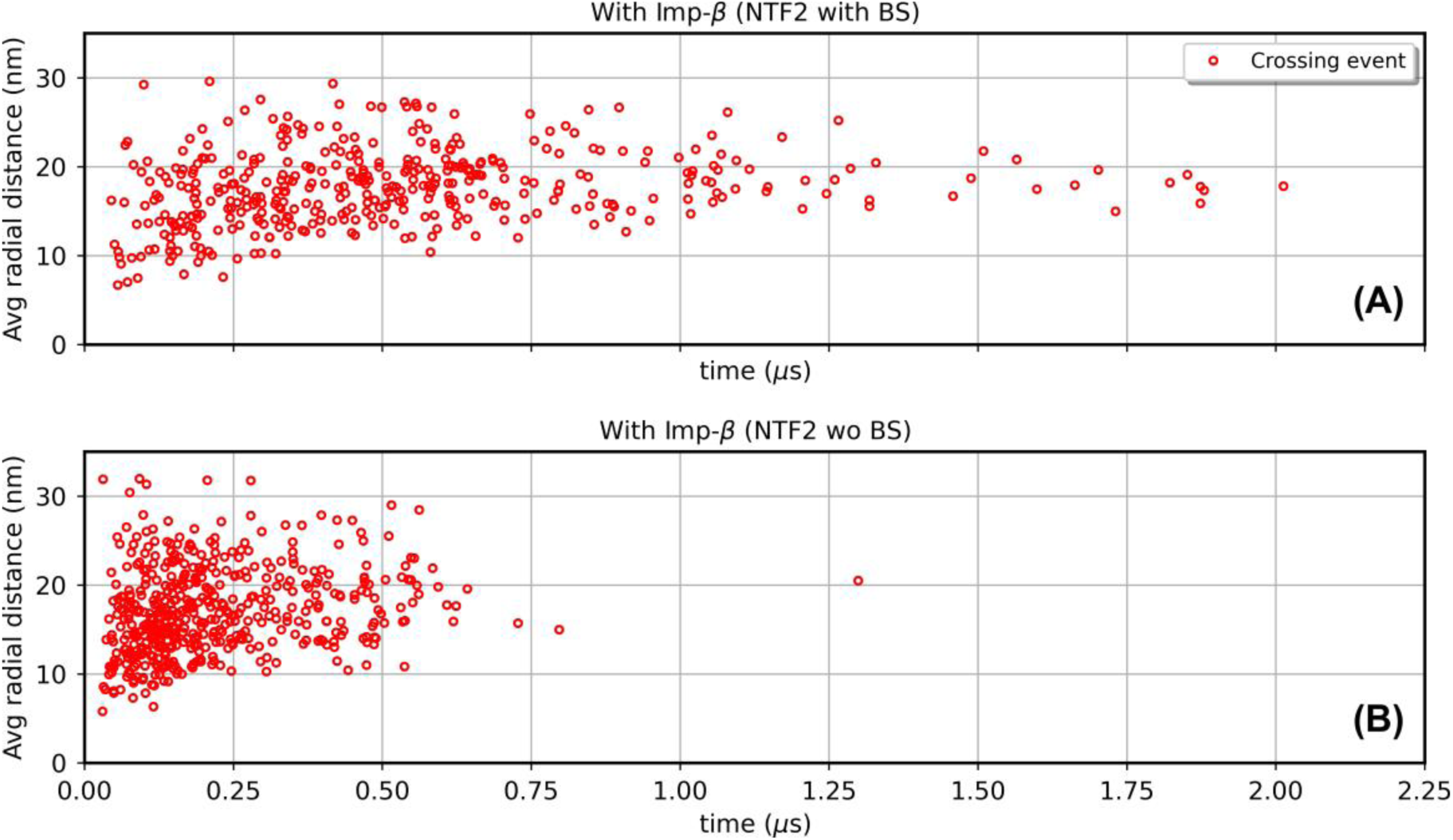
Average radial distance in the pore vs time it takes for the NTF2 to cross the pore for setup A (panel A) and setup C (panel B).

The observation that NTF2s move slower through the pore with hydrophobic binding sites can be explained by considering the role of specific interactions between NTF2s and FG-Nups.

In typical conditions, hydrophobic patches on NTF2s interact with FG motifs within the NPC, creating transient binding that can slow down their movement by tethering them to the FG-Nups. These interactions introduce a “stop-and-go” behavior, which aids in selective transport of large cargos but also increases transit time. In the absence of these hydrophobic binding sites, NTF2s are not subjected to this intermittent binding, allowing uncomplexed NTF2s to traverse the NPC more freely and quickly.

Although these results reveal key components of the interplay between FG-Nups and NTRs, it is important to note that the morphology of the nucleoporin chains depends on the force field chosen for this study, in particular, the choice of hydrophobic interaction parameters. In order to minimize error in predicted Stokes radii of FG-Nup segments vs experiments,^9^ Onck et al. made several changes to their original 1-bead per amino acid (1-BPA) force field.^30, 31^ The first change is the increase in hydrophobicity of charged residues as well as the addition of cation-pi interactions (incorporated into the 1-BPA-cp force field),^32, 40^ and the second is changing the hydrophobicity of highly abundant amino acids (G, Q and N) along with the exact masses of each amino acid i.e 1-BPA-1.1.^41^

For the work presented here, we initially tried 1-BPA-cp force field but this resulted in highly collapsed Nups which left a large empty space at the center of the pore. This was found to be a consequence of the increased hydrophobicity of charged residues which was initially introduced by Jafarinia et al. for the FG-Nup segments that contained more than 0.6% of Arginine content. In our setup the overall Arginine content is *∼* 0.009% (much lower than 0.6%). Therefore, we decided to implement the original 1-BPA force field with cation-pi interaction between F, Y or W with R or K.

Another important issue is the effect of (i) coarse-graining the protein components and (ii) adopting an implicit solvent model on the timescales observed in our simulations. This could explain the fast transit times (*µs*) observed in these simulations as compared to experimental values (milliseconds).^42, 43^ While previous computational studies report mean free passage times for molecules of similar size to Imp-*β* and NTF2 around 1–10 ms,^29^ we find faster transport. This may also be influenced by the high concentration gradient between the cytoplasm and nucleus, affected by recycling of transport proteins, which we adopted in order to enhance transit efficiency. Although absolute timescales from this work must be interpreted cautiously, the qualitative transport trends revealed in this work should provide valuable insights into nucleo-cytoplasmic transport dynamics. To our knowledge, this is the first computational study that elucidates the role of Kaps in regulating the transport of other NTRs through the NPC.

### NTF2 transport models

Analysis of our simulation results, achieved by radially dividing the pore into three distinct layers, reveals different transport patterns across all four setups. Figure 13 illustrates NTF2 transport patterns through each layer (shown in blue (Layer 1), orange (Layer 2) and black (Layer 3)) for all four setups.

**Figure 13:**
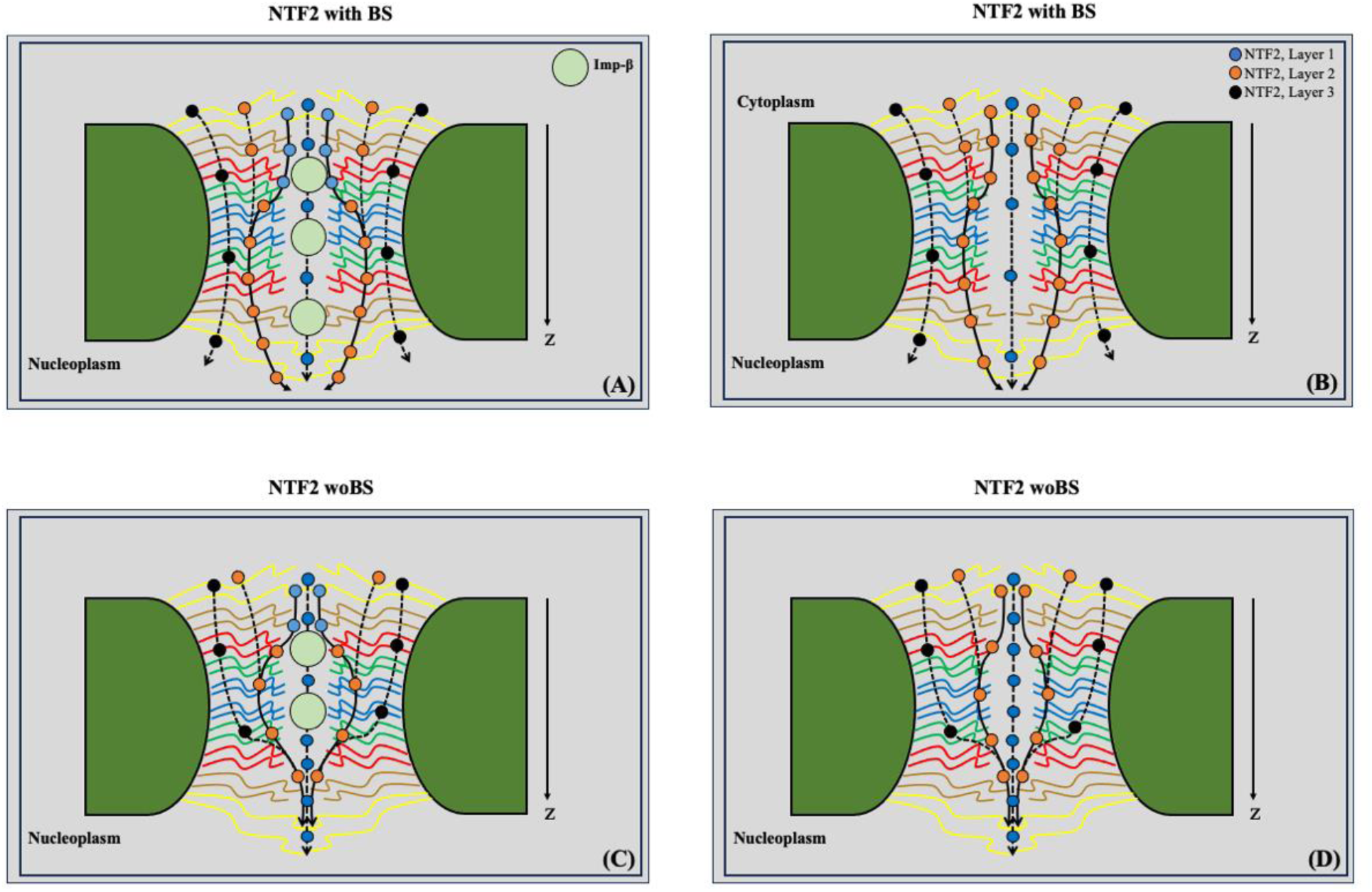
Transport patterns of NTF2s within the pore for all four steups. NTF2s passing through different pore layers are depicted using different colors: blue for Layer 1, orange for Layer 2, and black for Layer 3. Panel A vs. Panel B illustrates how, in Setup A, NTF2s are directed from Layer 1 to Layer 2 by Kaps and NTF2-F1G1 interaction. This directed transport is less pronounced in setup B, therefore resulting in a higher overall flux in setup A. Similarly, in setup C (panel C), Kaps initially push NTF2s into the pore, but these proteins exit through Layer 1 due to the absence of the NTF2-BS-F1G1 interaction.

The presence of Kaps appears to push NTF2 molecules into the FG mesh inside the pore. This interaction leads to a more concentrated flow of NTF2s through layer 2, resulting in a higher overall flux in setup A compared to B, where NTF2 do move into layer 2 due to their attractive interaction with the brush but their number is less due to a lack of a boost from Kaps. Furthermore, this boost by Kaps in setup A allows NTF2 molecules to penetrate deeper into Layer 2 (see Appendix, Fig. A4). Similarly, comparing panel 18C vs D highlights the effect of Kaps positioned in the center of the pore. In setup C, Kaps influence the flow by guiding NTF2s into region 2 at the pore entrance. However, due to the absence of attractive interactions between NTF2 and the brush, they do not linger in region 2; instead, they quickly transit and exit through region 1. This pattern of NTF2 flow is not observed in setup D. The impact of Kaps in redirecting NTF2 transport into layer 2, as observed in setups in which Kaps occupy the center of the pore, suggests a shift in the usual transport pattern. Consequently, this altered flow path in the presence of Kaps is likely responsible for the observed increase in overall flux in setup A vs B and setup C vs D.

## Discussion

This study highlights the crucial role of Kaps (Importin-*β*) in regulating NTF2 transport through the Nuclear Pore Complex (NPC), providing valuable insights into the “Kap-centric transport model”. Our simulations show that Kaps actively direct NTF2s into high FG-Nup density regions, expediting their transport and supporting the notion that Kaps not only serve as cargo carriers but can also regulate the transport of other NTRs like NTF2s. However, the exact mechanism of the Kaps affecting NTF2 transport, e.g., the sudden change in slope of the NTF2 transport count (Fig. 5) is still not clear and requires further investigation. Note that this change in slope was confirmed by a restart of simulation setup A at 4μs with a different random seed number for the Langevin dynamics, as shown in Appendix, Fig. A3.

Consistent with this Kap-mediated steering of NTF2s into FG-rich regions of the pore, recent experimental work has provided direct evidence for spatially organized transport pathways in the nuclear pore complex. In particular, Sau *et al.* used two-colour MINFLUX nanoscopy to visualize single-molecule trajectories of import and export receptors, revealing overlapping annular transport regions with reduced but nonzero occupancy near the pore center.^44^ These observations are consistent with our simulation results, which show preferential NTF2 transport through FG-rich annular regions (Layer 2) surrounding a lower-density core. In our model, interactions between transport factors and FG-Nups give rise to distinct transport pathways, with Kaps biasing NTF2 motion toward regions where passage is more efficient. Although the experimental study probes receptor dynamics *in vivo* while our approach relies on coarse-grained simulations, the qualitative agreement in spatial transport organization supports the relevance of the lanes model in NPC function.

In this context, our results are consistent with previous simulation studies that emphasize the role of FG-Nup density, crowding, and receptor occupancy in regulating nucleocytoplasmic transport. Prior computational work has shown that spatially heterogeneous FG-Nup organization can generate self-regulating transport behavior that prevents clogging and maintains efficient flux even at high NTR concentrations.^26, 27^ Our findings build on this framework by explicitly resolving how such density organization gives rise to spatially distinct transport pathways and lane-like behavior, particularly under Kap-rich conditions.^28^

We also note that our NPC model includes a reduced set of FG-Nups, following previous computational studies to maintain tractability. While this simplified architecture captures the overall physical environment of the central channel, additional FG-Nups such as Nup116, Nup100, and others may further influence mesh organization and receptor distribution. Moreover, the NPC is known to adopt multiple conformational states with different degrees of dilation, and variations in pore diameter are expected to modulate FG-Nup density and spatial organization, potentially reshaping the width, positioning, and stability of transport lanes. Incorporating additional Nups and exploring different dilation states in future work will enable a more complete assessment of how NPC composition and geometry shape transport pathways and receptor dynamics.

Finally, while our results reveal fundamental aspects of nucleocytoplasmic transport, they also highlight potential limitations due to the selected coarse-grained force field. The observed rapid transit times, influenced by the high concentration gradients, prompt further exploration into how model parameters affect absolute timescales. Future research integrating refined computational models and experimentalvalidation is essential to fully understand NPC dynamics and the complex role of Kaps in regulating selective transport. In summary, our results strongly support the Kap-centric transport model and demonstrate the existence of lanes that direct NPC transport.

## Acknowledgments

This work was supported by NSF grant CHE-1954865. The research was also supported in part by the University of Pittsburgh Center for Research Computing and Data, RRID:SCR_022735, through the resources provided. Specifically, this work used the H2P cluster, which is supported by NSF award number OAC-2117681. This work also used Bridges-2 at Pittsburgh Supercomputing Center through allocation sys160020p from the Advanced Cyberinfrastructure Coordination Ecosystem: Services & Support (ACCESS) program, which is supported by National Science Foundation grants #2138259, #2138286, #2138307, #2137603, and #2138296.^45^

## Data Availability

All simulation input files, force-field tables, and molecular data used in this study are publicly available at GitHub: https://github.com/skg43/KapCentric-NPC-Transport. The repository contains the exact LAMMPS input scripts and coarse-grained models used to generate the results reported in this manuscript.

## Appendix

### Binding sites in NTF2 and Importin-***β***

The binding sites on NTF2s were identified based on the work by Isgro and Schulten. Table A1 shows a list of residues these binding sites are composed of, while the residues shown in red in Fig. A1 have binding sites interaction with F1G1 residues in the Nups. The hydrophobic binding sites on Importin-*β* (Fig. A2) were similarly identified, and the residues that comprise these sites are listed in Table A2.

**Figure A1:**
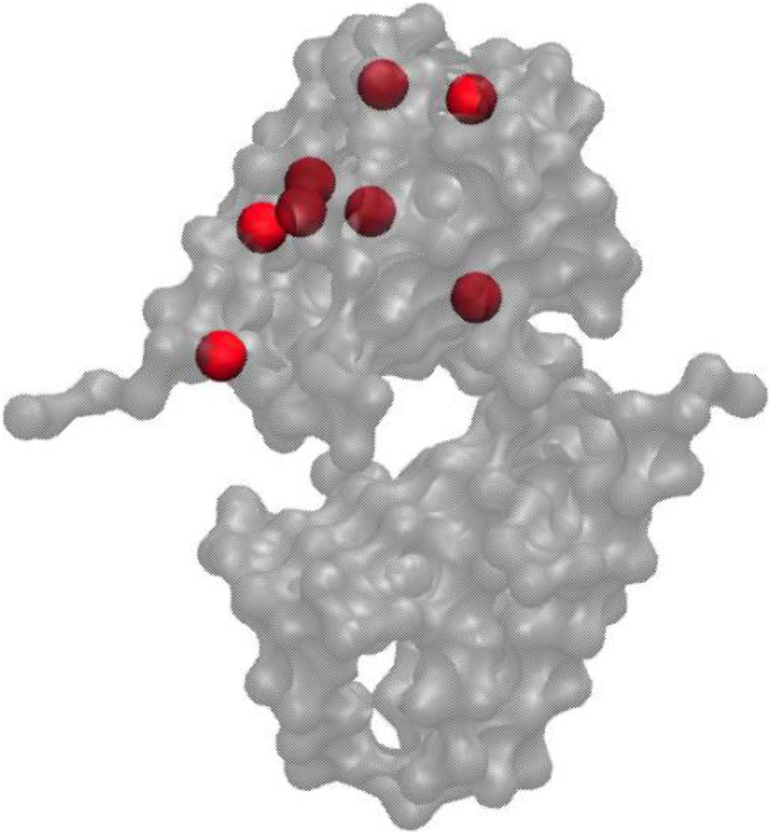
Binding sites on NTF2s identified based on the work by Isgro and Schulten.^35^

**Table A1:** List of binding site residues in NTF2 (PDB ID: 1OUN). Here bold residues are the ones with binding site interaction with F1G1 residues.

| Binding site | Residues |
| --- | --- |
| 1 | Ala35, Cys37, Gln46, Gly47, Asn115, **Trp6**, Pro74,<br>**Phe45**, Ile5, Trp6, Asp77, Ser78, Cys79, Ile80 |
| 2 | **Phe60**, Ile63, Leu88, Leu120, **Phe118**, Trp40, Leu58 |
| 3 | Ile9, Ser12, **Phe13**, His16, Ala110, **Trp111** |
| 4 | **Tyr18**, Gln15, Gln19, His65, Ile67 |
| 5 | Ile63, **Lys89**, **Lys62**, Asp91, Ala90, Gln64 |
| 6 | **Lys89**, Gln64, Ser66, Gln87, Ile95, His65 |

**Figure A2:**
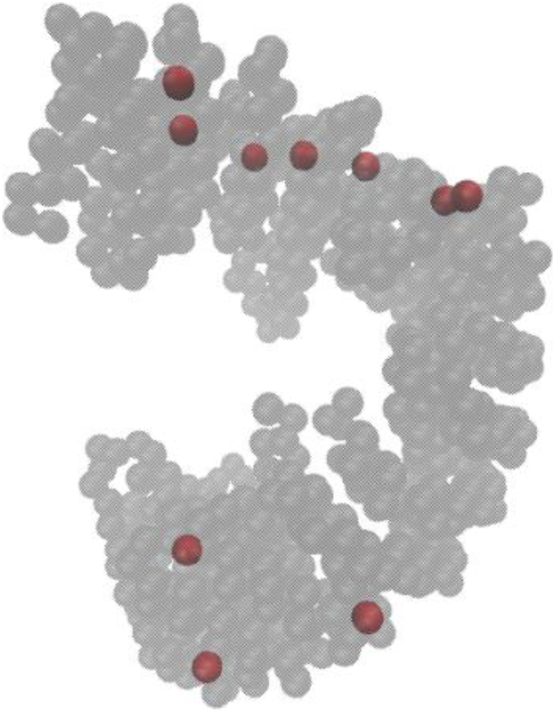
Binding sites on Importin-β identified by Isgro and Schulten.^34^

**Table A2:**
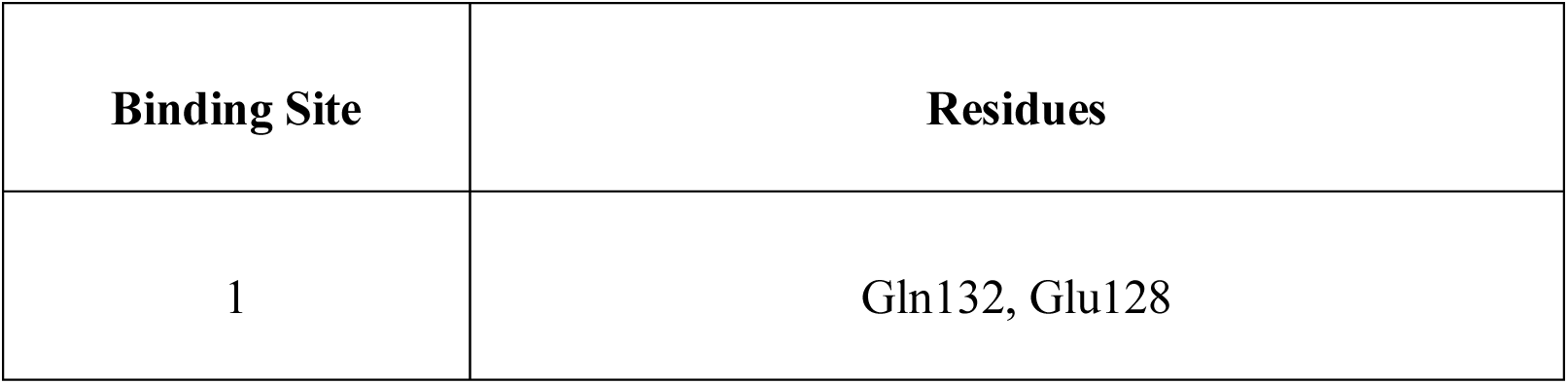
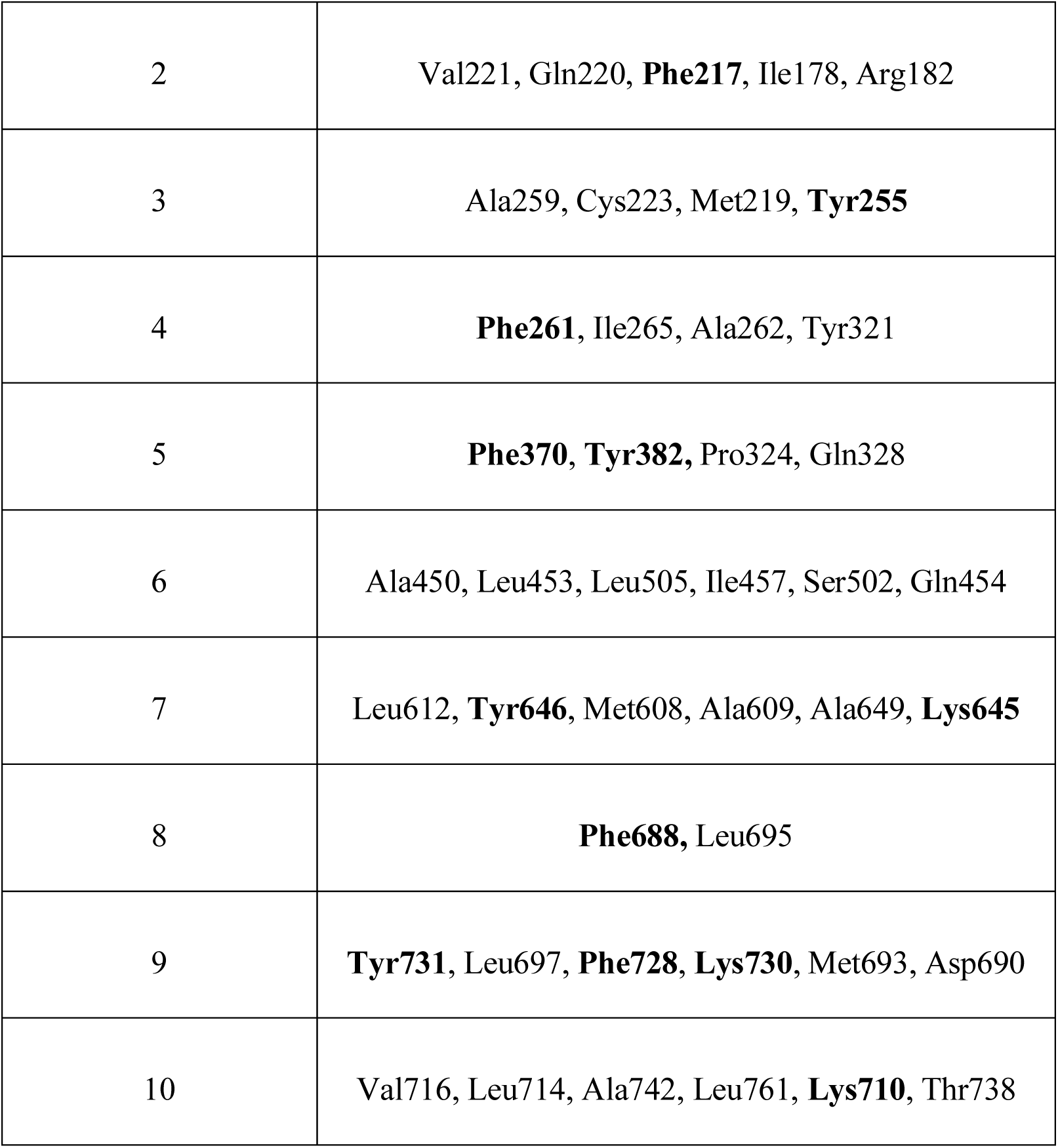
List of binding site residues in Importin-*β* (PDB ID: 2QNA). Here bold residues are the ones with binding site interaction with F1G1 residues.

#### Rerun of the simulation

**Figure A3:**
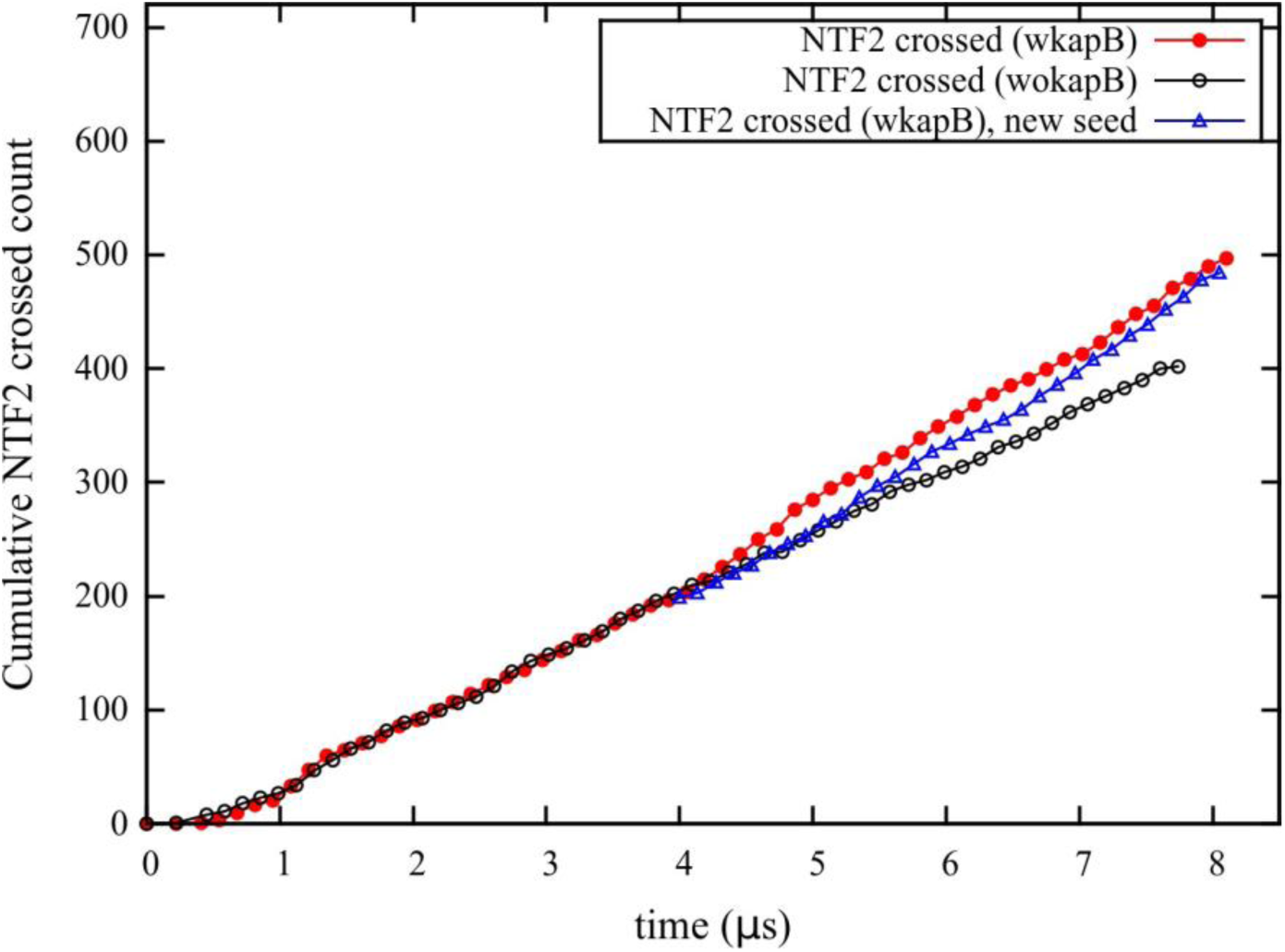
To confirm the slope change in the cumulative NTF2 crossing count in the presence of Kaps, a new simulation was run by restarting setup A at 4 *µ*s with a different random number seed (blue). We observed a change in slope similar to setup A which confirms the effect of Kaps on NTF2 flow.

#### Layer occupancy within the pore

**Figure A4:**
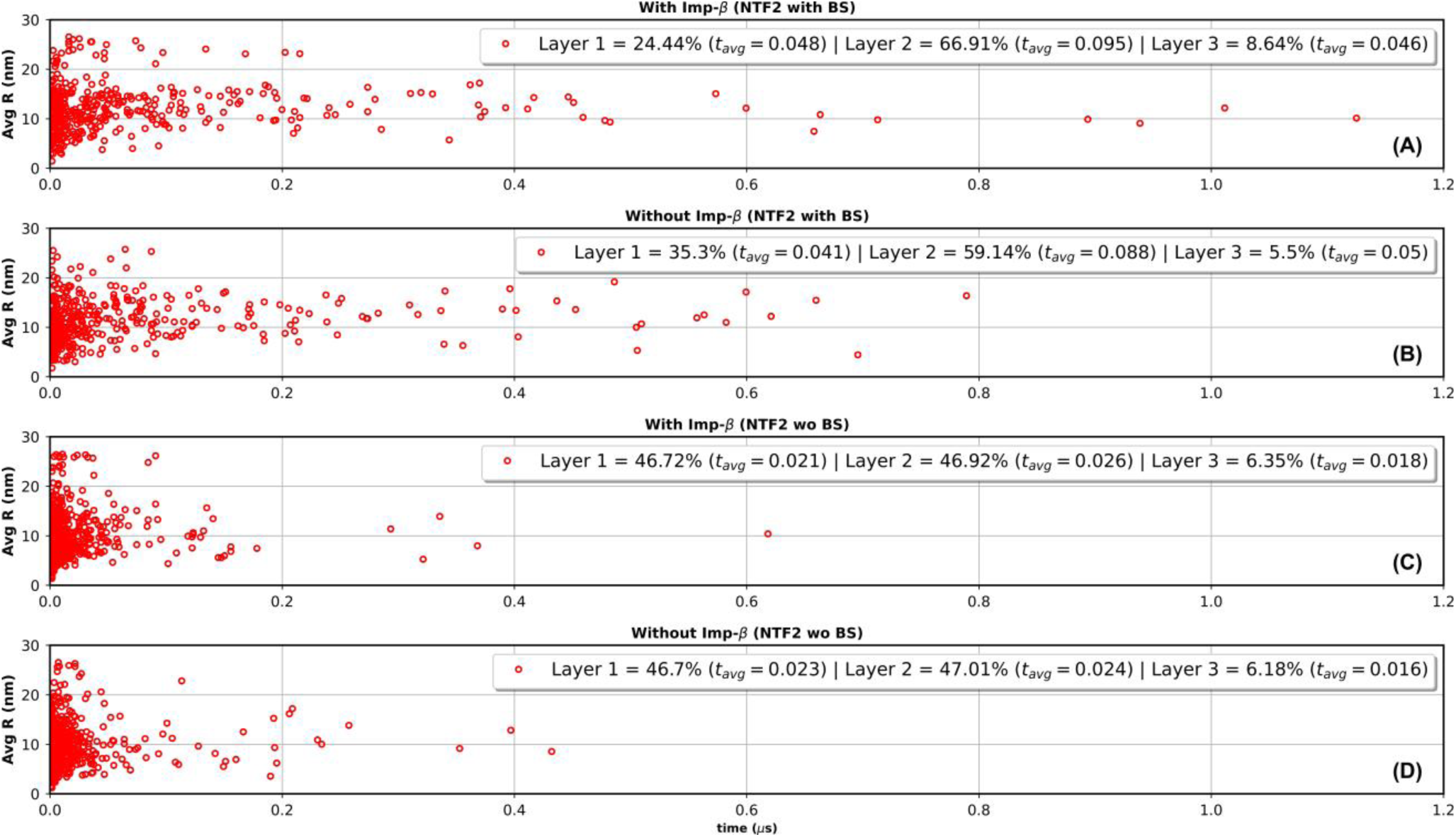
For each NTF2 that crossed the pore, its radial distance averaged over time that it spends in the central section (Avg R) vs time it takes to cross the central section is shown here. Additionally, the percent of crossing events through each radial layer and corresponding average crossing time (*t_avg_*(*µs*)) is shown for each setup. Notably, the percentage of NTF2s that crossed with an Avg R falling within Layer 2 and Layer 3 is higher for setup A (75.55%) vs setup B (64.64%). Therefore, NTF2s tend to traverse deeper into the FG mesh in presence of Kaps.

